# A Ray of Light Against Age Related Neurodegeneration

**DOI:** 10.1101/2023.04.25.538307

**Authors:** Elizabeth J. Fear, Frida H. Torkelsen, Kuan-Ju Chen, Martin Scott, Glenn Jeffery, Heidi Baseler, Aneurin J. Kennerley

## Abstract

Mitochondrial function declines with age and many pathological processes of neurodegenerative diseases stem from this dysfunction when mitochondria fail to produce the necessary energy required. Photobiomodulation (PBM), long-wavelength light therapy, has been shown to rescue mitochondrial function in animal models and improve human health, but clinical uptake is limited due to uncertainty around efficacy and the mechanisms responsible. Through combined theoretical Monte Carlo light modelling and practical ^31^P Magnetisation Transfer Magnetic Resonance Spectroscopy (MT-MRS) we quantify the effects of 670 nm PBM treatment on healthy aging brains. ^31^P MT-MRS revealed a significant increase in the rate of ATP flux after PBM in a sample of older adults. Light modelling shows 1% absorption in grey matter and confirms absorption peaks at 670 and 820 nm. Our study provides evidence of PBM therapeutic efficacy and strengthens confidence in PBM as an acceptable healthcare technology to improve mitochondrial function and human health.

## Introduction

Mitochondrial function declines with age due to time related oxidative damage, cysteine toxicity, mitochondrial DNA (mtDNA) mutations (either inherited or somatic)^1–4^ and impaired biogenesis.^5–10^ Neurological/psychological conditions (e.g., brain injury, stroke, Alzheimer’s disease, Parkinson’s disease, depression, anxiety and age-related cognitive decline) further render neuronal mitochondria vulnerable to oxidative stress.^11–13^ Alteration of the electron transport chain (ETC) ultimately reduces adenosine triphosphate (ATP) production and in turn increases apoptosis (figure 1a). Therefore, an aging population coupled with associated increases in cases of neurological conditions^14^ amplifies the need to develop safe, inexpensive treatments to restore mitochondrial function and offer neuronal protection as we grow old. Evidence shows that *non-invasive* transcranial red/infrared photobiomodulation (PBM) therapy can offer such neuroprotective benefits.^15, 16^

**Figure 1:**
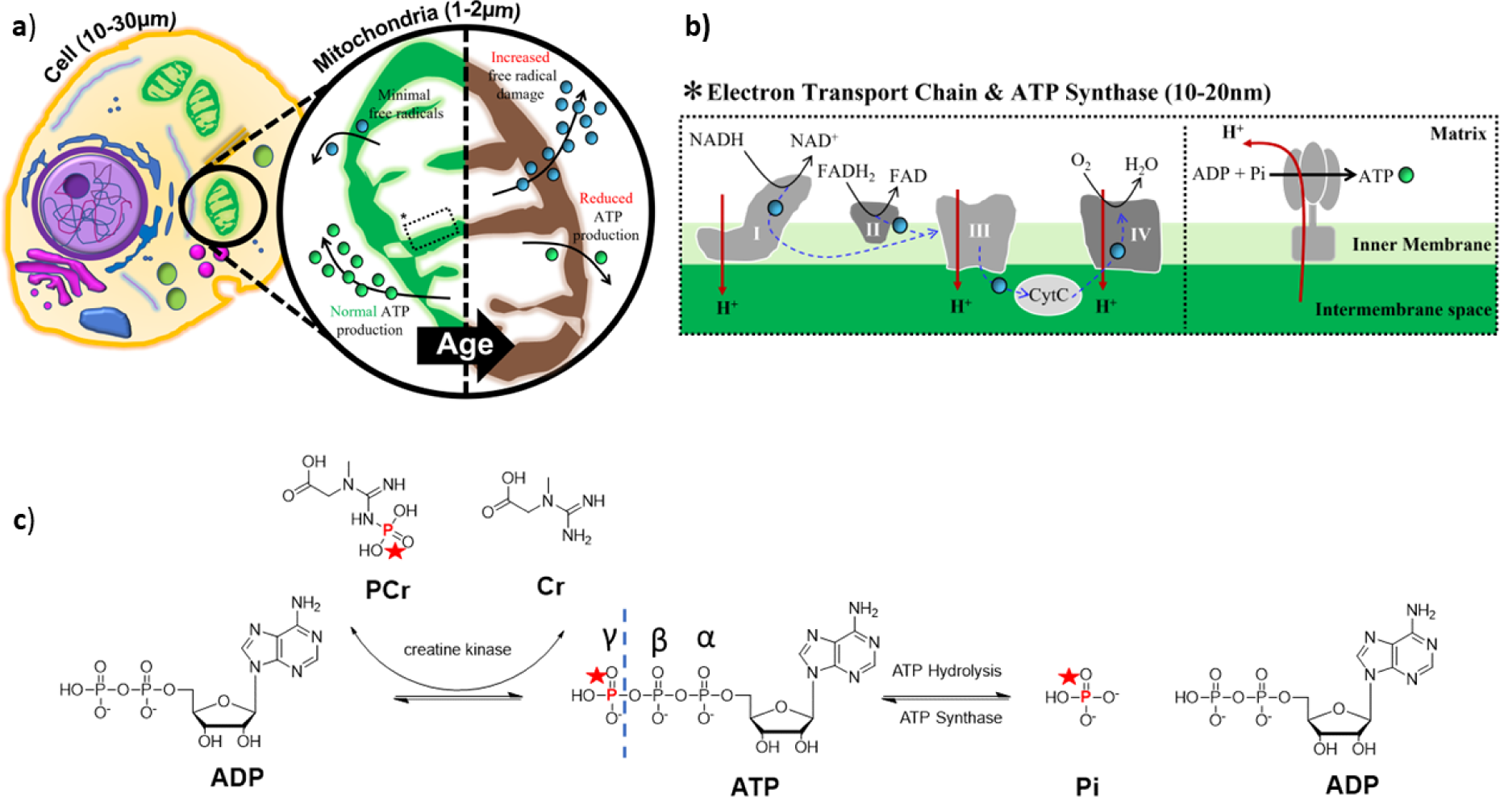
Key mediators of ATP production in the mammalian cell. a) Aerobic metabolism occurs within the mitochondria and drives ATP (·) production for normal cell function. With age, chronic exposure of mitochondrial membranes to free radicals/reactive oxygen species (ROS, ·), released a consequence of metabolism (or other), are hypothesised to drive decreases in cell function and viability.^1,2^ ATP production reduces, potentially accelerating age related neurodegenerative conditions;^3^ b) ROS are utilised as part of the electron transport chain (ETC) to alter membrane potential. A key mediator to help passage ROS through the various protein complexes (specifically III and IV) is cytochrome c. Movement of protons back into the mitochondrial matrix via ATP synthase produces ATP from ADP and inorganic phosphate (Pi). c) ATP is mainly generated through oxidative phosphorylation in the mitochondria, and to a lesser extent from glycolysis in the cytosol. Redox reactions, as part of the ETC, transfer electrons from Nicotinamide and Flavin adenine dinucleotide mediators (NADH & FADH_2_ respectively) to oxygen. Resultant energy is used to pump H^+^ from the mitochondrial matrix to the intermembrane space, creating an electrochemical proton gradient, which drives the synthesis of ATP from ADP and P_i_, via ATP synthase (figure 1b). ATP hydrolysis (i.e. use and breakdown back into ADP and P_i_) is high, even at resting state as ATP is required in many cellular processes. Creatine kinase (CK) in the cell cytoplasm also plays a role in maintaining the homeostasis of cellular energy. A reservoir of PCr is maintained at a concentration much higher than that of ATP. This makes it possible to carry out many energy-draining tasks quickly.

PBM illuminates the tissue with narrowband light in the 600 to 1100 nm wavelength range.^17, 18^ Reported benefits of PBM are wide reaching. In preclinical models PBM has been shown to improve inflammatory arthritis,^19^ aid wound healing,^20^ stimulate muscle regeneration^21^ and reduce necrosis in ischemic heart muscle.^22^ Improvements in sleep quality, mood and cognitive function have also been observed as a consequence of PBM therapy applied to the brain through the nasal cavity or via custom light helmets.^23, 24^ Recent studies have used PBM to reduce the lethality of COVID-19.^25, 26^

In terms of neuroprotection, transcranial PBM (usually in the near-infrared spectrum to penetrate through the scalp and skull) has been used to reduce damage caused by stroke (ischemic or other)^27, 28^ and remarkably has been shown to induce neurogenesis.^29–31^ Marked neurological improvements in Alzheimer’s disease,^32, 33^ Parkinson’s disease,^33–37^ depression and anxiety,^38^ ^39^ traumatic brain injury^40^ retinal aging and disease^41–45^ and general decelerated cognitive decline in aging^46^ have all been documented.

While data from animal models of ageing and neurodegenerative disease^41–44, 47^ have shown that PBM application improves mitochondrial membrane potential and therefore ATP production, clinical recognition of PBM therapy is limited by controversy over (i) the exact mechanism responsible for its positive effects; and (ii) poorly defined experimental design in terms of choice of wavelength, pulsed or continuous application, treatment time (duration and repetition), and even identification of the region of treatment application for particular disease conditions.

To promote clinical acceptance of PBM as a neuroprotective therapy, we present a proof of principle study utilising magnetisation transfer (MT) based ^31^P magnetic resonance spectroscopy (MRS) to quantify and track ATP production rate. We test the hypothesis that PBM therapy yields an increased rate of ATP production in healthy aging brains (60 years+).

ATP is mainly generated from adenosine diphosphate (*ADP*) and inorganic phosphate (*Pi*) via the aerobic pathway through oxidative phosphorylation (ADP + Pi → ATP) in the mitochondria via ATP synthase (figure 1b & 1c), and to a lesser extent via the anaerobic pathway from glycolysis in the cytosol. ATP hydrolysis (ATP → ADP + Pi) is high, even at resting state as ATP is required in many cellular processes.^48^ Hence ATP production during energy-draining tasks is supplemented through another anaerobic pathway, the creatine kinase (CK) pathway breaking down phosphocreatine, PCr, to creatine, Cr, in the cytoplasm (ADP + PCr ⇌ ATP + Cr) (figure 1c).^49^

Magnetic labelling of the terminal phosphate (γ) of ATP, with a radio frequency (RF) pulse train, induces a ^31^P MRS signal decrease in both Pi and PCr (through the two pathways described) – identifiable through differences in MR chemical shift. The signal reduction, Scr), for either Pi or PCr, as a function of saturation time, τ, follows an exponential decay:^50^

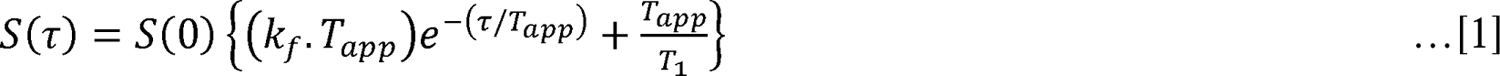

Where, k_f_ is the forward exchange rate towards ATP, *T*_1_ is the intrinsic relaxation time of PCr or Pi, and the ‘apparent’ time constant of decay, *T*_app_, is described as:

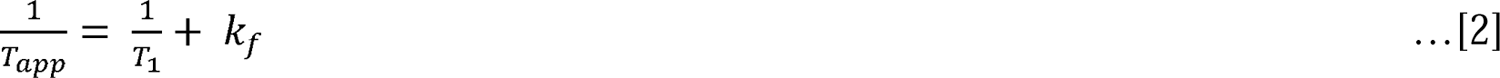

Therefore, by varying saturation time and measuring signal levels, one can estimate the forward exchange rate of ATP for both pathways using equations 1 & 2.

Here we present a refined non-invasive *in-vivo* imaging protocol that can be used in future studies of disease and neuroprotection with PBM. Our protocol is supported with extensive theoretical exploration of light transport through tissue. We used Monte Carlo approaches to simulate light absorption and scattering within layered tissue models of the human head. The tissue models deployed were informed by structural MRI data from individual participants to help elucidate potential mechanisms of PBM action.

The leading mechanism of PBM therapeutics involve mitochondria homeostasis. Within mitochondria the balance between reactive oxygen species (ROS) and ATP production is preserved by integrating multiple cellular signals.^51, 52^ When too many ROS are produced, major disturbances in cell function and viability occur, leading to disease (figure 1a).^52–55^

The inner mitochondrial membrane houses carrier proteins (complexes I-IV) responsible for the induction of an electrochemical proton gradient to drive ATP synthase (as part of the ETC – figure 1b). The enzyme cytochrome C oxidase (CCO), known as complex IV, catalyses the rate determining step in the ETC.^56^ CCO is also involved in the formation of apoptosome and therefore the progression of apoptosis.^55^ CCO is a photoreceptor with a broad absorption spectra (figure 1c). Within oxidised CCO the haem iron-Met80 bond causes increased light absorption at ∼695nm (figure 1d).^55^ The copper centres in CCO have a broader absorption range that peaks around 830 nm (figure 1e).^57^ It was therefore believed that photoexcitation of the electronic states of these compounds changes the redox properties, fundamental for effective generation of the electrochemical membrane proton gradient driving ATP synthase as part of the ETC.^58^ Indeed, Wong-Riley et al. (2004) illuminated primary neurons (with 670, 728, 770, 830, and 880 nm light), functionally inactivated by toxins.^59^ The greatest increase in energy metabolism resulted under 830 nm and 670 nm light. The least effective wavelength was found to be 728 nm. This was attributed as evidence for up-regulation of CCO. An array of *in vitro* cell culture data now exists confirming increases in ATP synthesis following PBM.^60–62^ Furthermore, any absence of these near-infrared absorption bands implies a dysfunctional/denatured conformation of CCO and is often referred to as an ‘indicator of trouble’. Keeping CCO upregulated with photoexcitation was hypothesised to inhibit release into cytoplasm and stave off cell death/aging.

However, maximum light absorbance for CCO occurs <425 nm.^63^ Therefore, it would be expected that ATP production rate would be superior, illuminating cells at this wavelength of light. Counterintuitively, when human adipose stem cells were exposed to blue light (415 nm, 16 mW cm^-2^) a reduction of intracellular ATP levels was observed (with an increase in intracellular ROS). Longer exposure to this blue light resulted in further decreases in ATP levels.^47, 63, 64^ Alternative mechanisms involving a reduction in interfacial water layer (IWL) viscosity in the irradiated cells were therefore proposed.^65, 66^

The beneficial effects of infrared based PBM could also be caused by photo based generation of singlet oxygen, localized transient heating of absorbing chromophores, and increased superoxide anion production with subsequent increase in concentration of the product of its dismutation, H_2_O_2_.^58^ Nitric oxide (NO) is proposed to be released upon cell illumination (photo-dissociation of NO from the binuclear centre (a_3_/Cu_B_) of CCO)^17^ which causes vasodilation and therefore increased blood flow to the vicinity of the cell. Whatever the direct mechanism, these effects have been reported to be short-term i.e. occurring during PBM treatment.^15, 67^ ^68, 69^

Formation of ROS after CCO excitation has also been suggested as a mediator of the beneficial biological effects of PBM (780 nm illumination) therapy.^70^ ROS are important in the activation of transcription factors in the nucleus. These transcription factors lead to the up-regulation of stimulatory and protective genes leading to cell proliferation/neurogenesis.^71^ Importantly such effects have been found to continue after PBM treatment and may explain the positive effects found days, weeks and months after treatment.^15, 67, 68^

While the exact mechanism of action for PBM remains contested, this study is the first to probe the metabolic benefits of PBM therapeutics in-vivo in the ageing human brain. Note the longitudinal relaxation time, *T*_1_, of Pi and PCr are longer than the lifetime of ATP^72, 73^ at clinically relevant MR field strengths. Key reaction kinetics relating to ATP production can therefore be probed non-invasively with magnetisation transfer (MT) Magnetic Resonance Spectroscopy (MRS).^74^ Here we use ^31^P MT MRS to measure ATP metabolism in aging brain pre and post PBM application.^72, 75, 76^ Experimental data is supplemented with theoretical exploration of light transport through tissue to better inform future PBM therapeutics.

## Materials and Methods

### Participant information

Written informed consent for this study was obtained from all participants. Ethical approval was granted by York Neuroimaging Centre (YNiC) Research, Ethics and Governance Committee. This study followed the tenets of the Declaration of Helsinki. Ten healthy participants aged 60+ were recruited from the YNiC volunteer list (mean age = 68 years; age range = 60-85 years; 6 females, sex assigned at birth). General observations regarding hair colour, coverage and skin tone (all participants were white British) informed light transport and statistical models. Volunteers were excluded from the study if they had contraindications for completing the MRI procedures, known neurodegenerative conditions, were currently enrolled on an interventional clinical trial, and/or if they were unable to comply with the study.

### Experimental Design

A 5-day longitudinal study was designed (figure 2a). Baseline ^31^P MT-MRS assessments were completed on the morning of day 1. Immediately after scanning, participants underwent their first session of PBM and training for self-application for the next three days. PBM was applied before midday over a four day period in accordance with data from Weinrich et.al.^77^ & Shinhmar et.al.^78^ On day 5 participants completed a further ^31^P MT-MRS assessment to investigate changes in ATP flux. We constructed this 5-day experimental design based upon Wong-Riley et al. (2005) who showed that LED treatment at 670 nm significantly reversed the detrimental effect of a toxin on neurons using a similar cumulative dosage (p<0.001 for three metabolic neuronal types).^59^

**Figure 2:**
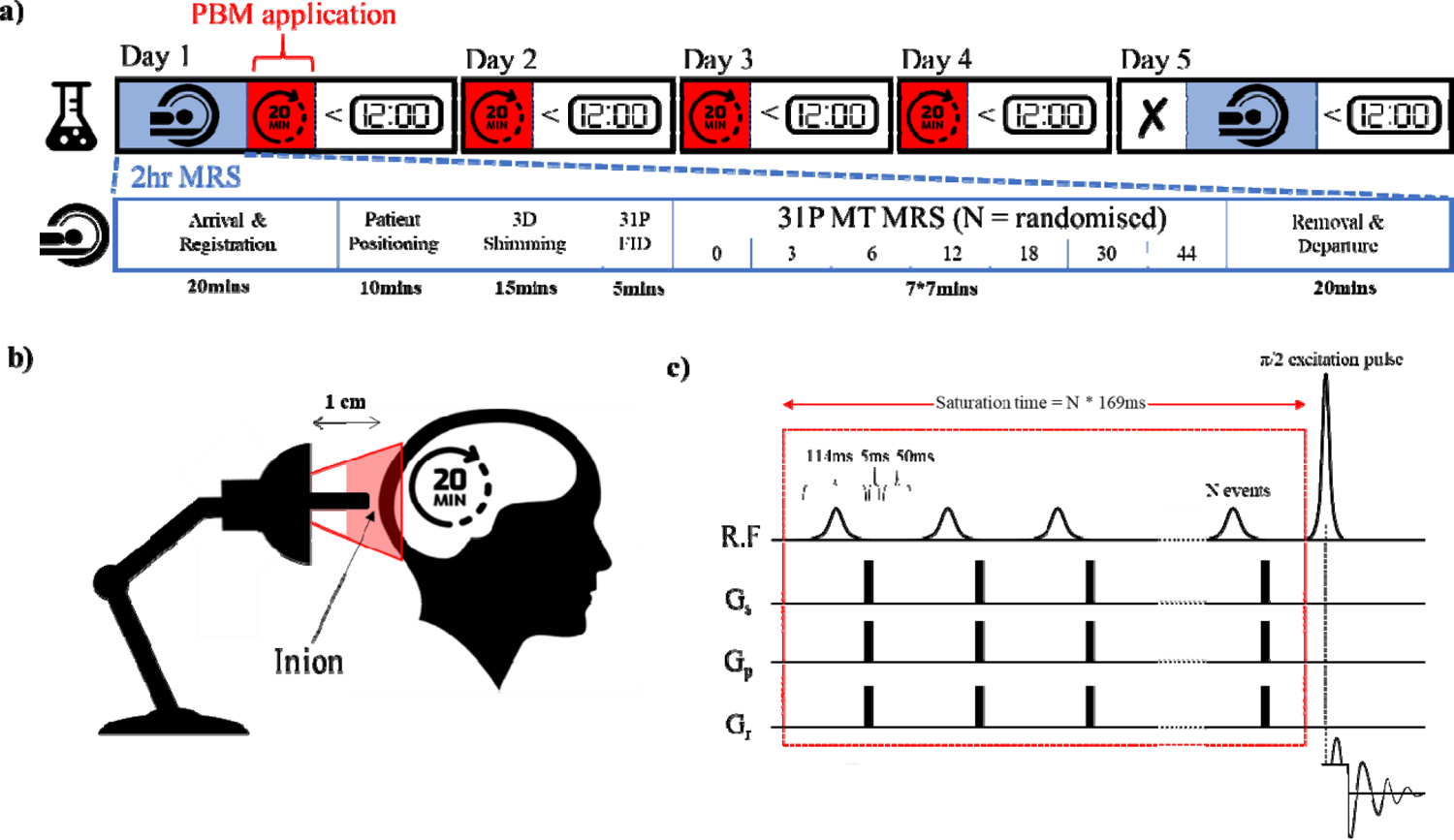
Experimental design for tracking the neuroprotective effects of PBM treatment. a) Baseline ^31^P MRS measurements were taken on all participants on day 1 followed by the first session of PBM for 20 minutes applied to the back of the cranium This was repeated for the next three days before midday. On day 5 participants returned for follow-up MRS measurements (N.B. no PBM was applied on this day); b) PBM therapy utilised a 670 nm LED bulb positioned approximately 1 cm from the inion; c) ^31^P MRS utilised a magnetisation transfer FID based pulse sequence with a varying number of 114 ms saturation RF pulses (0, 3, 6, 12, 18, 30 and 44) followed by a 5 ms spoiler gradient and 50 ms delay. Experiment order was randomised for each scanning session.

PBM treatment used a narrowband (peaking at 670 nm) E27 Edison screw (27 mm diameter) flat ended light bulb consisting of 12 LED sources with a 30-degree beam angle (18W, Red Mini 670, Red Light Man Ltd., Manchester, UK), mounted on a spring-loaded (angle-poise) adjustable arm. This allowed the participant easy placement while sat/positioned comfortably. The centre of the bulb was held 1 cm from the participant’s inion using a small spacer (figure2b). PBM therapy targeted the occipital lobes of the human brain. The comparatively high concentration of metabolites found in the brain, which are often altered in neurodegenerative diseases, can be exploited.^79^ The occipital lobes were selected because of their ease of access, density of grey matter and proximity to the MRS surface coil used for measurement in the scanner. The light was administered for 20 mins (participants were supplied with a digital timer). Participants were instructed to sit as still as possible during PBM application, but to immediately stop application if the scalp felt uncomfortably warm.

### Magnetic Resonance Spectroscopy

MRS measurements utilised a 3T MAGNETOM Prisma system (Siemens Healthcare GmbH, Erlangen, D). A dual channel ^1^H/^31^P transmit-receive flexible surface coil (O-XL-HL-030-01150 V03; Rapid Biomedical GmbH, Rimpar, D) was used for RF transmission/detection. The coil was positioned at the head of the scanner bed and participants placed in a supine position. The inion was positioned over the centre of the flex surface coil and foam cushioning used to limit lateral bending and aid comfort. Magnet safe padding was used to curve the flexible coil on either side of the head. Participants were instructed to stay as still as possible during measurements. No head restraints were used, and participants were allowed to listen to music while in the scanner. Participants were given foam ear plugs to protect hearing during scans.

Following positioning of the head at the magnetic isocentre of the scanner, a ^1^H based *T*_2_ localizer was acquired (3 slice packages; 15 slices; 256 mm^2^ Field of View (FOV); 6 mm slice thickness; Repetition Time (TR) 606 ms; Echo Time (TE) 122 ms; Number of averages (NA) 1, Flip Angle (FA) 150°; 60% phase resolution; 6/8 phase partial Fourier; 592Hz/px; echo spacing 4.06 ms; turbo factor 154; RF pulse type – fast; gradient mode – fast). This tri-planar localiser was used to position a cuboid adjustment window covering the entire visual cortex (figure 3a). Manual frequency, power and 3D shimming adjustments were completed to achieve a ^1^H water peak with Full Width at Half Maximum (FWHM) of 19.9 ± 1.0 Hz and ^31^P PCr peak with FWHM of 16.5 ± 0.97. 3D shimming utilised a multiple gradient echo field mapping approach with resolution optimised for brain imaging. The PCr resonance was set to 0 ppm. This was confirmed by a short ^31^P FID based acquisition (TR 4000 ms; NA 32; FA 90^°^; bandwidth (BW) 4000 Hz, acquisition duration 512 ms; spectral points 2048) further used to find the exact frequency offset for γ-ATP.

**Figure 3:**
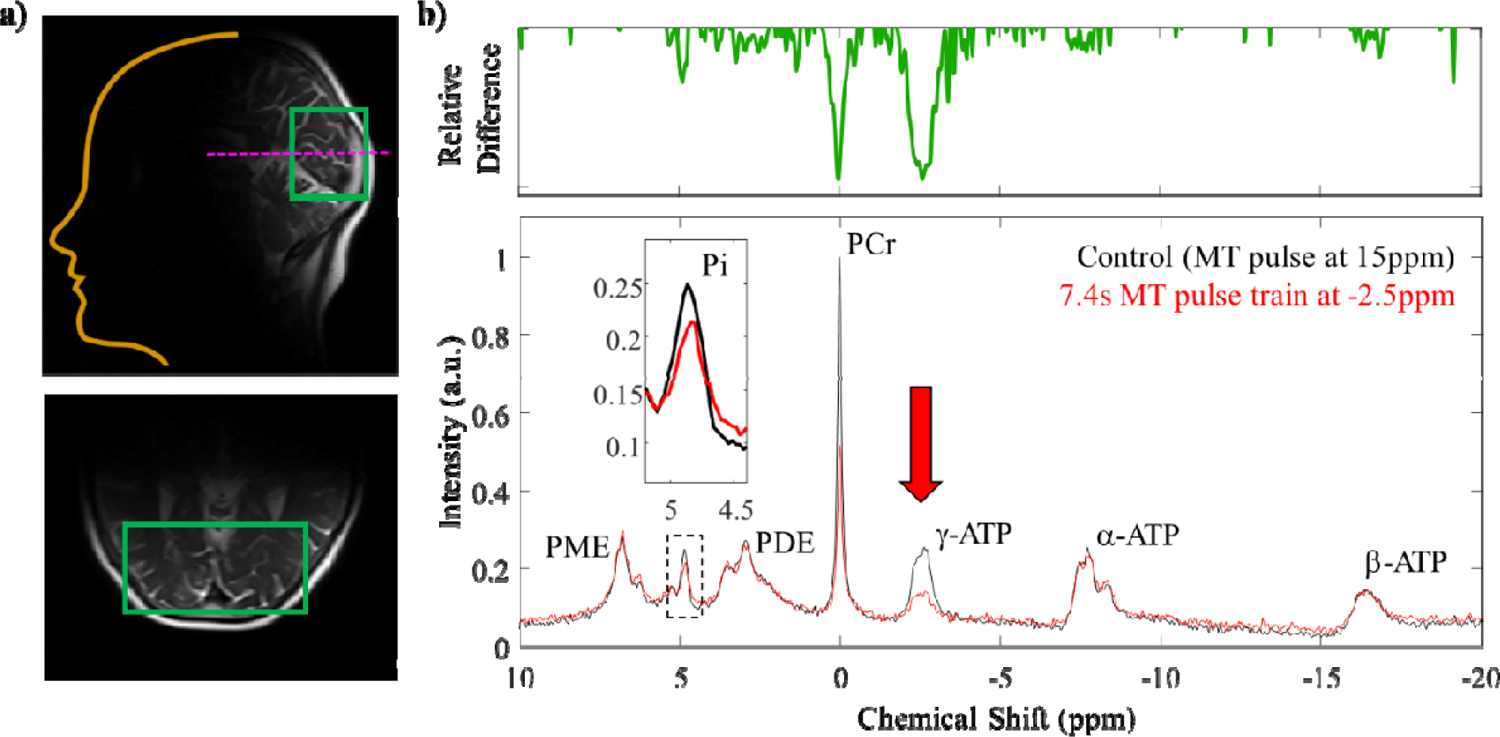
Representative ^31^P MR spectra. a) A standard T_2_ localiser was used to position a cuboid adjustment window (green box) covering the entire visual cortex. Following suitable frequency matching, power calibration and shimming adjustments, ^31^P spectra were acquired. b) Phased and baseline corrected ^31^P spectra (40 ppm spectral width) were used to identify key metabolites via chemical shifts for the three phosphorus sites of adenosine triphosphate (ATP – gamma, γ *@-2.5ppm, alpha, α @-7.5ppm and beta, β @ −16ppm), phosphocreatine (PCr @ 0ppm) and inorganic phosphate (Pi – intracellular (i) @ ∼4.9ppm and extracellular (e) @ ∼5.2ppm). Spectral peaks for phospho-mono- and di-esters were also observed. For tracking ATP flux rates multiple amplitude*L*modulated (selective) RF pulses of constant maximum amplitude and length (2.06*Lμ*T, 114.29 ms) were applied on the γ-ATP resonance. This saturated the γ-ATP signal. Chemical exchange of this phosphorus nuclei resulted in observable drops in signal for PCr (creatine kinase pathway) and Pi (ATP synthase pathway). Note – control experiments applied 30*MT pulses at 15 ppm to account for saturation of the underlying phospholipids*.

A non-localised ^31^P free induction decay-based magnetisation/saturation transfer (MT) sequence with narrowband selective saturation was developed in-house (figure 2c) following Chen et.al.^80^ The MT sequence pulse train was comprised of multiple amplitude[modulated RF pulses (hyperbolic secant in shape) of constant maximum amplitude and length (2.06 µT, 114.29 ms) interleaved with spoiler gradients (5 ms, 10 mT/m) in G_x_, G_y_ and G_z_ and a 50 ms delay time. Following the MT pulse train, a standard pulse-acquire excitation regime (nominal FA 90^°^) was applied.

The MT RF pulse allowed sufficient irradiation of the ^31^P γ-ATP resonance (@ approx. − 2.5ppm) but selective enough to have negligible sidebands. To assess the selectivity of the MT pulse, ^31^P spectra were acquired across 3 experiments with 30 pulses applied off centre at i) −2.5 ppm (the intended γ-ATP resonance), ii) +2.5 ppm (the chemical shift of phosphodiesters, PDE) and iii) +15 ppm (off-resonance). As expected, (due to limited chemical exchange between the target species) no drop in signal for Pi (@ 5ppm) or PCr (@ 0ppm) was observed when the MT pulse was applied at +2.5 ppm, confirming suitable selectivity. Off-resonance saturation with 30 pulses at +15 ppm was used as the baseline condition (0 s saturation time). Phospholipid molecular rotation is fast compared to the saturation time, resulting in a broad resonance under the peaks of interest.^81^ ^82^ RF saturation at −2.5ppm will therefore affect both phospholipids and γ-ATP. Saturation at 15 ppm effects only phospholipids resulting in comparable baselines.^83^

The time permitted for chemical exchange between ATP and PCr or ATP and Pi (or saturation transfer time, t_sat_) was varied across seven separate scans by changing the number of RF pulses in the MT train. Number of pulses was varied as 0, 3, 6, 12, 18, 30, 44 x 169 ms (RF pulse 114 ms + delay 50 ms + spoiler gradients 5 ms), resulting in saturation transfer times (t_sat_) of ∼0 s, 0.5 s, 1.0 s, 2.0 s, 3.0 s, 5.0 s and 7.4 s. The order of the seven scans was randomised for each participant at each visit. A pause of 60 seconds was applied between each MT scan to allow for complete spin relaxation to equilibrium. Other scan parameters were as follows: NA= 32; TR = 12000 ms; BW = 3000 Hz; Data points = 2048; acquisition duration 692 ms. The total scan time, including setup, was approximately 80 minutes.

### Spectral analysis

The ^31^P spectral data were analysed offline in MATLAB 2020a (The MathWorks, Natick) using software routines developed in-house. Raw frequency data underwent 5Hz line broadening (Gaussian filter) to improve SNR. Spectra were manually phased (0^th^ and 1^st^ order correction). Following Fourier transform the baseline was fitted to a 4^th^ order polynomial and removed. Resultant spectral peaks were assigned to the following ^31^P resonances: phosphomonoesters (PME), Pi - intracellular ∼4.9 ppm and extracellular ∼5.3 ppm), phosphodiesters (PDE), PCr and the three resonances of adenosine triphosphate (γ-ATP, α-ATP, and β-ATP) (figure 3b).

To quantify signal contributions from intracellular (Pi), the spectra were further windowed between 4-6 ppm. Data were fitted, using nonlinear least squares with a Levenberg-Marquardt algorithm, to appropriate Gaussian/Lorentzian functions for Pi(i) and Pi(e) with chemical shift, amplitude and FWHM floating variables. The baseline (raised due to phospholipid contributions) was again fitted to a fourth order polynomial. Fitting parameters were used to isolate the signal contribution from intracellular Pi. The area under the curve was extracted as a function of saturation time for the seven scans. All data were normalised to the control (15 ppm offset) experiment.

The formation of ATP via the ATP synthase pathway (aerobic) and the creatine kinase pathway (anaerobic) are in equilibrium (figure 1). Therefore, signal saturation of the terminal phosphate (γ) of ATP, with a pulse train of RF, induces a signal decrease in both PCr (due to the CK equilibrium) and P_i_ (due to oxidative phosphorylation). Resultant MT ^31^P-MRS data were fitted to equation 1 and 2. We followed Chen *et. al.* and used a fixed 3.0s for the *T*_1_ of Pi(i), to extract the rate of ATP flux, *k_f_* (analysis was repeated for PCr with an assumed *T*_1_ of 5.1s).^80^

### Statistics

Due to the relatively small sample size and non-normally distributed data (Kolmogorov-Smirnov test of normality, D = 0.262, *p* = 0.010), non-parametric statistics (Wilcoxon Signed Ranks test, two-tailed) were applied to compare pre- and post-PBM *k_f_* rates for both metabolic pathways (Pi and PCr). As participants varied in age (60-85 years), a Kendall’s tau (non-parametric) correlation was performed between age (years) and the difference in k-rate before and after PBM treatment to assess whether age played a significant role in PBM effects.

### Light Transport through Tissue Modelling

We modified a 3D Monte Carlo Simulation (MCS) developed by Jacques et al. (2013)^84^ and original based upon a simple 1D approach ^85^ to investigate light transport through tissue layers (defined by cubes each denoted by simple integers to identify tissue type e.g. skin, skull, brain etc.). Our model incorporates key chromophores including water, melanin, cytochrome-c and haemoglobin. The model was used to calculate illumination levels required to mitigate melanin absorption and quantify photon weight reaching grey matter. This allowed us to explore whether absorption by cytochrome-c is a possible mechanism of PBM action.

Although simple 2D layered tissue models (and validated against Wang et.al. 1995) support a nuanced assignment of tissue properties in three-dimensional space, such models often assume matched boundaries, with tissues sharing refractive indices, and do not account for boundary surface orientation. Tissues have varying thickness in different regions across 3D space. Layers or slabs do not accurately represent the curved or uneven surfaces of many tissues. Boundary curvature would influence the photon trajectories, as the angle of a tissue boundary relative to the photon direction influences the likelihood of transmission. In the case of the grey matter surface, the presence of sulci and gyri will affect the paths of photons to different extents depending upon the photon entry point.

Here, our model used a tissue model constructed from triangular meshes of the tissue/volume surface. These meshes were generated from previously acquired T1 and T2-weighted structural MRI data where available (3 of 10 participants). FreeSurfer (Martinos Center for Biomedical Imaging) was used to generate a ‘brain mask’ and segment grey and white matter volumes (GM & WM respectively). In MATLAB the brain mask was inverted and applied such that simple intensity filtering on the image data could generate voxel masks for the skin, skull and cerebral spinal fluid (CSF) (figure 4a).

**Figure 4:**
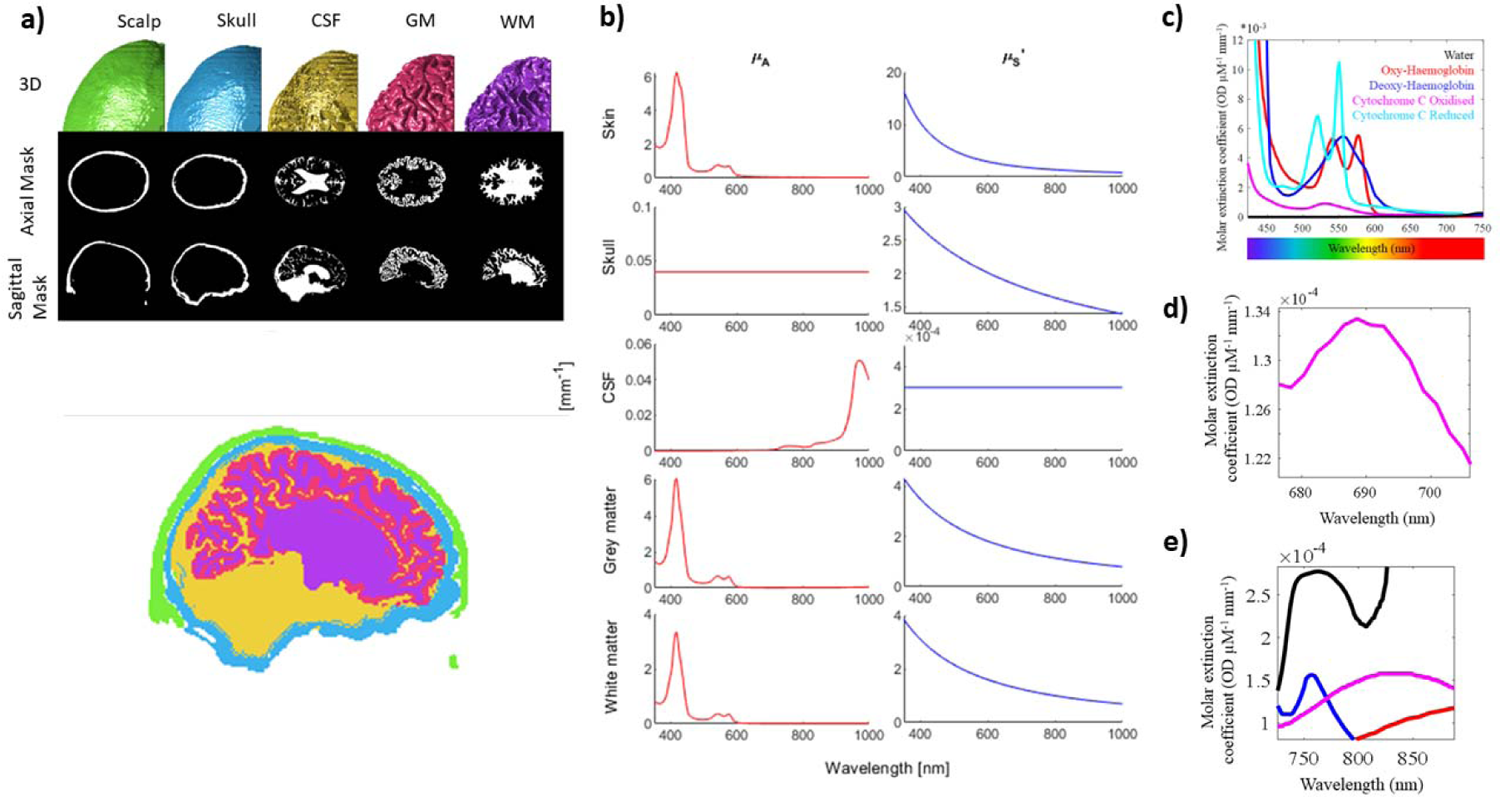
Tissue model for Monte Carlo Simulation of Light Transport through tissue. a) High resolution T1 and T2 MR image data was used to segment scalp, skull, CSF, GM and WM tissue structures to inform our 3D MCS. b) optical properties for each layer across the illumination wavelengths were determined from existing literature data and equations 3-10. c) Cytochrome is a key chromophore in the brain with molar extinction coefficients similar to haemoglobin; d) The oxidised state of cytochrome c has a small peak in absorption at 695nm due to the haem iron-Met80 bond; e) The copper centres in cytochrome c oxidase also have a broad absorption range that peaks around 830 nm. These photon wavelengths are in the near infra-red range and able to penetrate the skin and skull to reach brain tissue for activation of cytochrome c via absorption.

A 3D mesh was generated based on the voxel data of each mask using the Marching Cubes algorithm. Originally presented by Lorensen and Cline in 1987 for visualising MRI and CT data,^86^ the method has undergone development, notably by Chernyaev^87^ to improve topology preservation, and is today considered robust and computationally cheap, making it the most popular algorithm for extracting iso-surfaces from volumetric data.^88^

The trigonometric calculations involved in the MCS were updated to account for the complex tissue boundary orientations extracted above. Our updated MCS included i) collision tests for photons intersecting with tissue boundaries (following the Moller-Trumbore algorithm^89^); ii) calculation of photon direction after reflection or refraction with an angled boundary surface; and iii) indexing of absorption events by their complex tissue ‘layer’ defined by explicit x, y, z coordinates. Simulations covered the photon wavelength range 350-1000 nm, in increments of 5 nm in order to assess the most appropriate illumination wavelength for maximum cytochrome absorption. The mesh-based simulations were tested with a simple one-layer tissue model with the following optical properties: refractive index n = 1, μ_A_= 10cm^−1^, μ_S_= 90cm^−1^, anisotropy factor g= 0.75 and thickness 0.02cm. Ten simulations were completed each one following 50,000 photons through the model. Predictions of diffuse reflectance (R_d_ = 0.09761) and total transmittance (T_t_ = 0.66112) (both unit less quantities) were validated against Wang et al. (1995) and demonstrated excellent agreement with this accepted method (R_d_ = 0.09734 ± 0.00035; T_t_ = 0.66096 ± 0.00020).

Light transport through tissue is heavily dependent upon the scattering and absorption properties of the turbid media through which it passes. Jacques (2013) provides an overview of tissue optical properties as determined across the literature.^90, 91^ Based upon phantom work by Wrobel et.al. ^92^ and a survey by Lister et.al.^93^ our simulation used a constant 1.365 refractive index for all tissue types. The anisotropy factors, g, were similar to previous models^92^ (0.88, 0.94, 0.999, 0.96 & 0.87 for skin, skull, CSF, GM & WM respectively). We combined Mie and Rayleigh scattering effects to estimate the reduced scattering coefficient, µ’_S_, of skin (equation 3).^94^

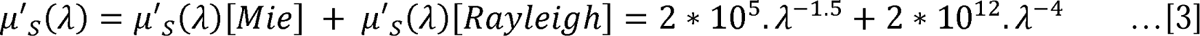

The μ*’_S_* for CSF was set to a constant 3·10^−4^ mm^−1^ across all wavelengths. Studies have indicated values between 0 and 0.3 mm^−1^ which will not significantly affect light transport pathlength estimates.^95^ The scattering properties of skull, GM and WM were calculated using equation 4 following Jacques,^90, 91^ The data were converted into fitting parameters according to this equation, where the wavelength (λ) was normalized by a reference wavelength (500 nm). Therefore parameter ‘a’ is the value of μ*’s* at 500 nm and was used as a simple scaling factor. The value *b* is referred to as the scattering power, resulting in a wavelength dependence of μ*’_S_*. We used listed typical values for a and b values for bone, brain (GM) and WM.^94^

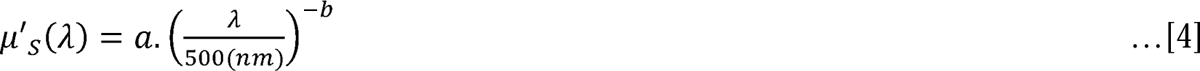

Wavelength dependent absorption coefficient, μ*_A_,* values for skin were estimated ^94^. Mean scalp thickness from MRI data was 7.4 ± 0.9 mm. Epidermal thickness (*d_epi_*) was assumed to be 0.1 mm of this scalp depth (*d_skin_*), the ratio of which was used to weight the final estimated skin absorption coefficient (equation 5). The model included both epidermal (equation 6) and dermal (equation 7) absorption. The epidermal absorption coefficient was calculated from melanin absorption (equation 8) and a skin baseline absorption (equation 9) coefficient. All absorption coefficients were calculated in mm^-1^ units. The melanin fraction, *f*_mel_, was set to 0.04, representing a typical value for the light-skin tones (and matching the ethnicity of the participants in the present study). Blood fraction in skin (*b_f_*), typically between 2-5% in well-perfused tissue, was set to 0.03 (i.e. 3%).^94^

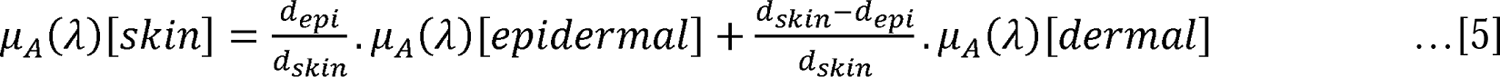

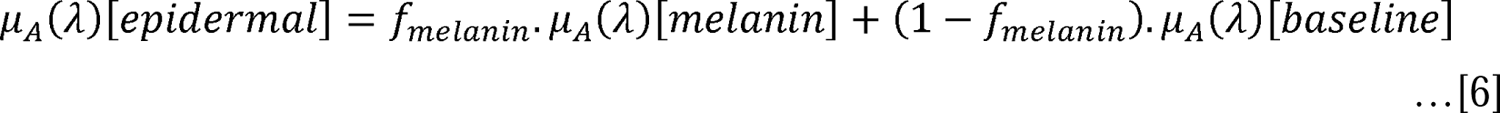

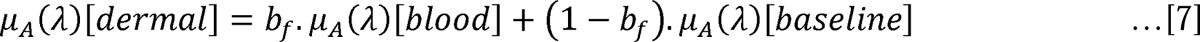

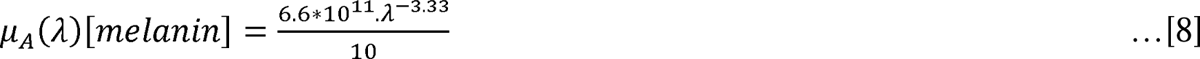

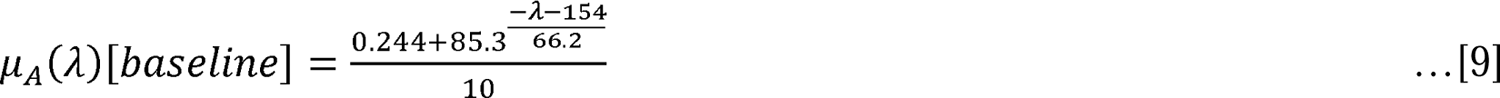

The μ*_A_* value for skull was set to a constant 0.04 mm^−1^ across all wavelengths. CSF absorption was assumed to be similar to that of water. GM and WM μ*_A_* values were estimated from extinction coefficients, ε_i_ (figure 4c-e), and assumed concentrations, [c_i_] of the five main chromophores (oxyhemoglobin (HbO_2_), deoxyhemoglobin (Hbr), cytochrome-C oxidase (CytC-ox), cytochrome-C reduced (CytC-red) and water) using equation 10.

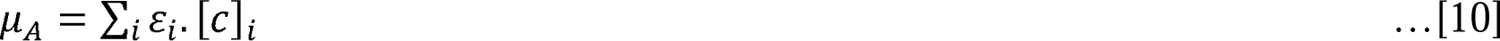

The assumed concentrations of the chromophores are listed in Table 1. These are based on previously estimated concentrations,^96^ but adjusted to fit the water and blood content difference between white and grey matter.^97^. Final wavelength dependent optical properties for each segmented tissue are shown in figure 4b.

**Table 1:**
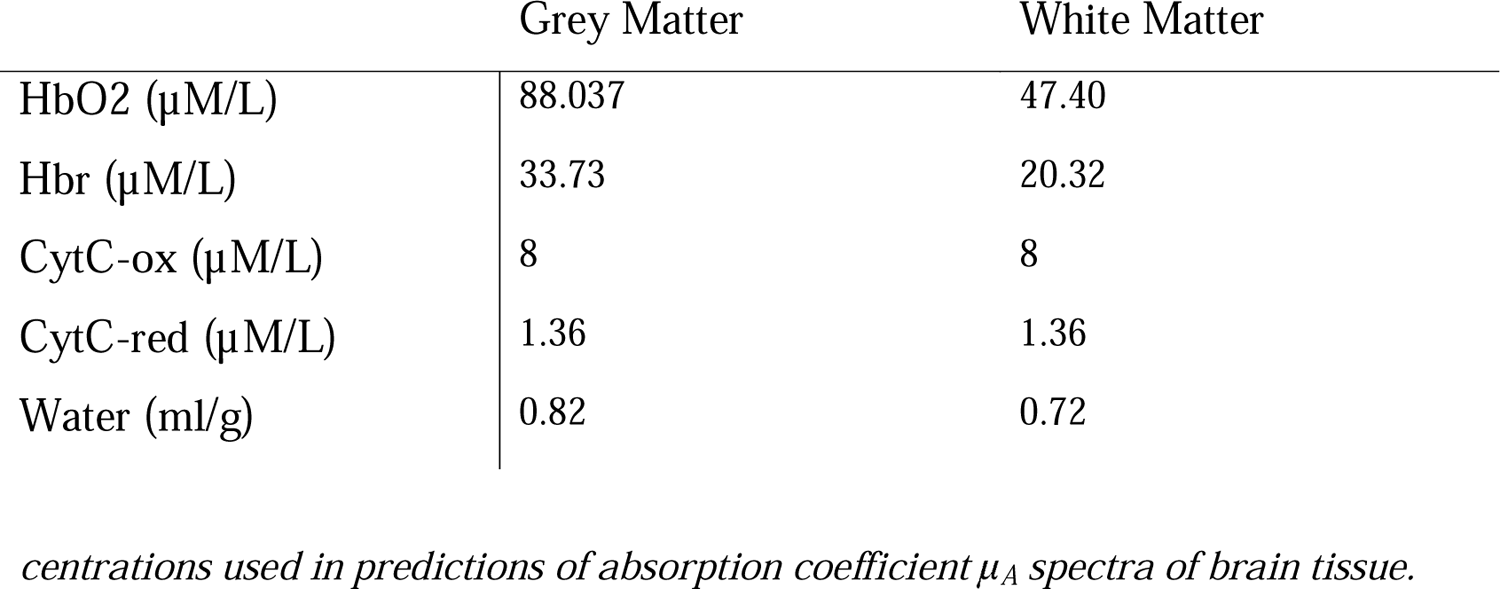
Chromo phorecon

Rather than running one long simulation (>10 h simulation/job for 50,000 photons), the workload was divided into 100 smaller simulations (nodes) of 500 photons each running on a high-performance computer cluster. Lower absorption coefficient values within some layers extended job runtimes. In those cases, photon number was limited to reduce computation time. Developed MATLAB code is available online at: https://github.com/fht502MCS

Note, skull and CSF layer thickness are known to be dependent upon brain region, age and gender.^98^ ^99^ Simulations were completed where mean skull and CSF thickness were inflated to explore these physiological confounds.

## Results

### General Observations

Over the 5-day treatment period, none of the 10 participants reported uncomfortable/adverse heating of the scalp due to PBM application over the 20-minute application time confirming that 1 cm is a suitable distance between PBM light source and head to maximise photon count. The ^31^P MT pulse sequence developed remained within safe Specific Absorption Rate (SAR) constraints for MRI and again no participant reported feeling hot or uncomfortable during the 50-minute ^31^P MT MRS measurement time.

However, of the 10 participants who completed the study, three were excluded from analysis. One participant self-reported low-level migraine each morning that were present on day 1 of the study (baseline MRS data collection) but had dissipated by day 5 (follow-up MRS data collection). While the participant anecdotally declared this a ‘miracle’ of PBM, we excluded this participant because the pathophysiology of migraine in terms of vascular contributions is still debated.^100^ Furthermore, changes in cerebral blood flow and potential cortical spreading depression events^101^ associated with migraine ^102, 103^ can drive energetic imbalance. A second participant, while asked to remain still within the scanner, had an involuntary movement condition. This resulted in increased noise in the spectral measurements which made it difficult to identify a suitable Pi peak. A third participant required a comfort break during the baseline (day 1) MRS measurement and had to be removed from the scanner. Two points on the MT decay curve were measured before the break, five after, confounding analysis.

Due to on-going COVID-19 disruptions, two of the remaining 7 participants could not return on day 5 of the experiment. Treatment was extended to 7 days, and these two participants returned for ^31^P MRS assessment on day 8. This is noted in the subsequent analysis.

### ^31^P Magnetisation Transfer Spectroscopy

Representative ^31^P spectra from the human visual cortex are shown in figure 3. Key metabolites were easily identifiable. When applying the magnetisation transfer (MT) pulse train (30 pulses) off resonance at 15 ppm, we observed clear peaks for the three phosphorus sites of adenosine triphosphate (ATP – gamma, γ @-2.5 ppm, alpha, α @-7.5 ppm and beta, β @ −16 ppm), PCr @ 0ppm and Pi – intracellular (i) @ ∼4.9ppm and extracellular (e) @ ∼5.2ppm). Spectral peaks for phospho mono- and di-esters were also observed.

With the MT pulse train (44 pulses – longest saturation time of 7.4 s) off resonance at −2.5ppm (chemical shift of γ-ATP) we see comparable spectra to that acquired with saturation at +15[ppm. This is direct evidence that similar suppression of the phospholipid baseline was achieved between the conditions. While the off[resonance 5 s saturation time (30 pulses in the MT train) provides adequate phospholipid suppression, we did not explore if this could be achieved with smaller saturation times.

Importantly, in the ^31^P MT spectra we see a clear/effective saturation of the target γ-ATP peak. Figure 3b shows the relative difference between this spectrum and the above ‘control’ condition. We also observe drops in peak amplitude for Pi and PCr due to chemical exchange as part of aerobic and anaerobic metabolic process outlined in the introduction. We note that while the associated drop in Pi signal amplitude is small, it is easily detectable by eye at the long saturation times used in this experiment (see figure 3b insert).

The chemical shift difference between PCr and γ[ATP is only ∼2.51Lppm, and partial suppression of PCr while saturating γ[ATP has been documented.^104^ Hence, in an additional control experiment we moved the centre frequency of the saturation pulses to 2.5ppm to quantify any partial suppression effects. By following Chen et.al. (2017) and using adiabatic hyperbolic secant pulses, we see negligible direct saturation due to RF bleed (rather than chemical exchange) when using these highly selective pulses (data not shown). No further corrections to the data were applied.

Representative results for progressive saturation of γATP are shown in figure 5 a-d. Associated signal decreases in Pi(i), due to reversible chemical exchange as part of ATP synthase, were visible in the ‘raw’ filtered spectral data at long saturation times (7.4s – highlighted in figure 5a). However, the inherent low SNR for this important metabolite (driven by the clinical field strength used (3T) and the short acquisition period – 6.4 minutes), meant accurate data quantification of peak area required data fitting (see methods). ^31^P MT based spectra (windowed between 4 and 6 ppm) were fitted to two Lorentzian functions corresponding to Pi(i) and Pi(e). Chemical shift, peak amplitude and peak FWHM were floating variables. Resulting fits are shown in figure 4b. Again, the drop in Pi(i) signal with increased saturation time is maintained and clearly observable. No similar trend was observed for Pi(e). This could reflect the lack of chemical exchange pathways between γ[ATP and Pi(e) or the even lower SNR due to low tissue concentration. It is noted that there was still a significant baseline offset – even following phospholipid saturation. Therefore, a fourth order polynomial was fitted to the data allowing extraction of the peak of interest – in this case intracellular Pi (figure 5c). The area under the curve was extracted and plotted as a function of saturation time (figure 5d). For each saturation time, error was estimated as the RMS error from the fitting residual. An exponential decrease in signal was observed as a function of saturation time. This was fitted to equation 1 to estimate the ATP flux rate (*k_f_*) (figure 5e).

**Figure 5:**
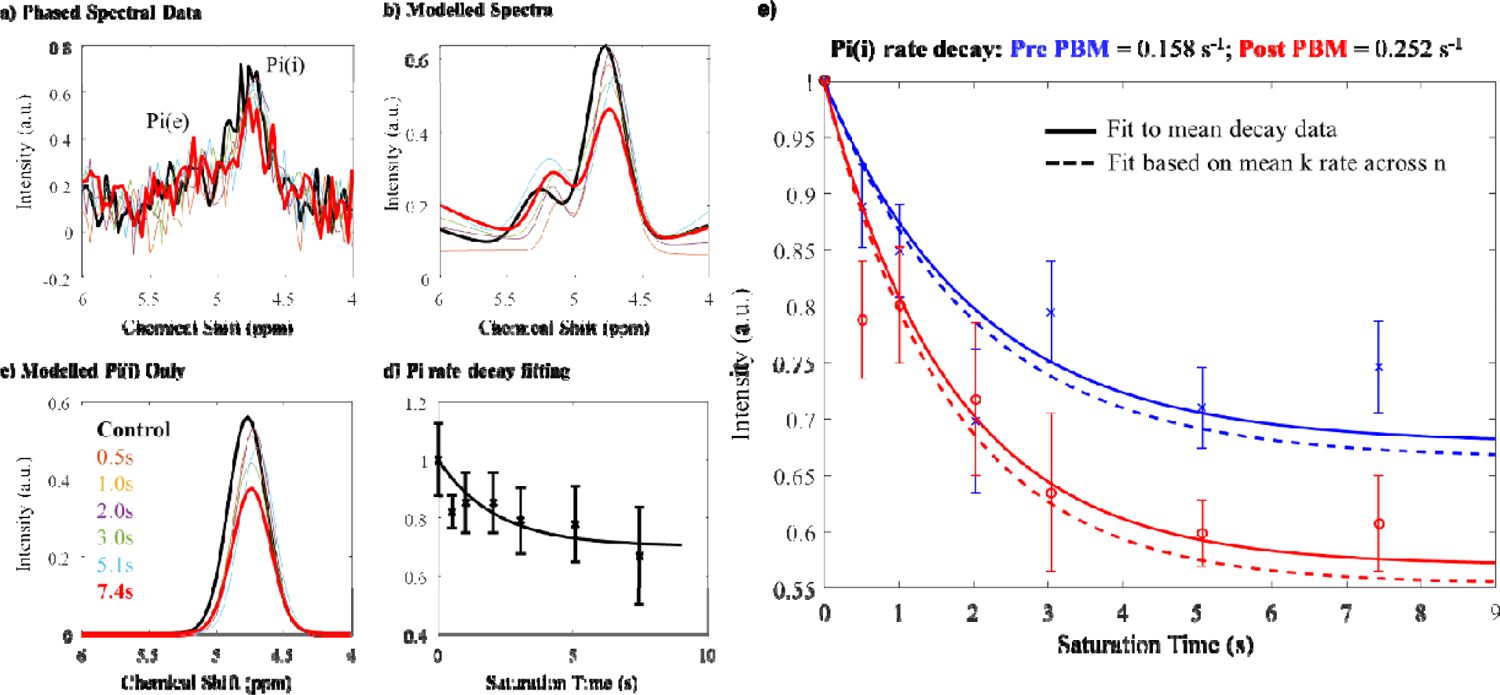
Data analysis pipeline for MT-^31^P MRS data. a) Raw FID data was filtered using an exponential windowing function and phased before Fourier transform into spectral space for baseline correction. Data were windowed between 4-6 ppm to isolate the Pi peaks; b) Resultant spectra were fitted to appropriate Lorentzian functions for Pi(i) and Pi(e) with chemical shift, amplitude and FWHM free variables. The baseline was fitted to a fourth order polynomial; c) Signal contributions from only Pi(i) were isolated for each saturation duration and area under the curve taken and normalised to the control (no saturation) experiment; d) subsequent data were plotted against saturation time and the resulting decay curve fitted to estimate ATP flux rate (equation 1). e) ATP flux rate estimates pre (blue) and post (blue) PBM treatment based on signal decay as a function of RF saturation time. We find an increase in mean k rate for ATP production post PBM treatment. T_1_ of Pi(i) was fixed to a constant (3.1s) pre- and post-treatment. Curves were fitted for each individual participant and also for the mean decay data across participants. Standard errors across n=7 participants shown.

Note that while these data can be used to fit both forward ATP flux (*k_f_*) and *T*_1_ for Pi(i), with only 7 data points in the extracted exponential decay, and relatively high noise for each point, floating both variables caused the fitting algorithm to hit the boundary conditions for *T*_1_(Pi(i)) in all cases (1 or 7s). We therefore followed Chen et.al.^80^ and fixed *T*_1_ at 3.0s and only extracted *k_f_* for each individual participant (n =7) and for the mean decay data (see figure 5c).

Fitting to the mean decay curve across n=7 participants pre- and post-PBM application, we found a *k_f_* rate for ATP synthase of 0.158 s^-1^ and 0.252 s^-1^ respectively. Fitting each individual decay curve and then averaging extracted *k_f_* rates, we found 0.198 ± 0.035 s^-1^ and 0.298 ± 0.045 s^-1^ pre- and post-PBM respectively. We found an increase in *k_f_* rate, for ATP production post-PBM compared to pre-PBM application. For transparency, when data from the three excluded participants (see above) are included, extracted *k_f_* becomes 0.182 ± 0.025 s^-1^ and 0.208 ± 0.044 s^-1^ pre- and post-PBM respectively.

Change in ATP flux pre- and post-PBM was evaluated statistically. Figure 6a-c summarises the changes in the rate of decay of Pi before and after PBM treatment. The rate increased in six of the seven participants and remained stable in one participant. Because of the small sample size and non-normally distributed data (Kolmogorov-Smirnov test of normality, D = 0.262, p = 0.010), non-parametric statistics were applied to compare pre- and post-PBM rates. The decay rate for Pi (intracellular) was significantly faster post-PBM compared to pre-PBM (Wilcoxon Signed Ranks test, two-tailed, Z = −2.366, p = 0.016).

**Figure 6:**
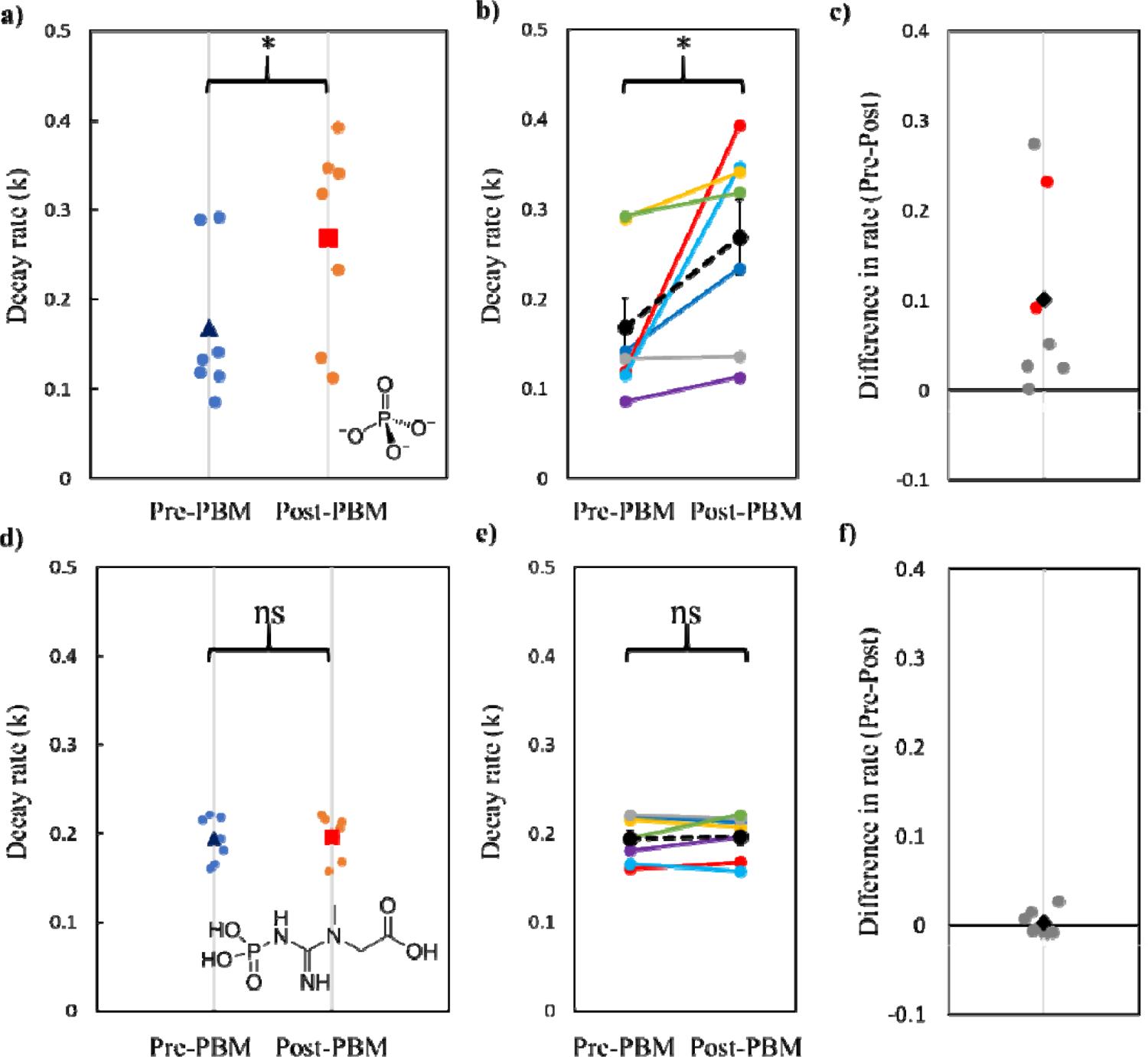
Exploration of key rates of decay before and after PBM treatment. a) Relative decay rate (k) of inorganic phosphate pre- and post-PBM treatment. Small dots represent individual data. Blue triangle: mean k rate pre-PBM. Orange square: mean k rate post-PBM. A significant difference was evident (Kolmogorov-Smirnov test of normality, D = 0.262, p = 0.010); b) Relative decay rate (k) of inorganic phosphate pre- and post-PBM. Coloured lines indicate individual data pairs (n=7). Dashed black line and dots represent the mean of all participants; c) Difference in k-rate (post-PBM – pre-PBM). Small dots indicate individual data. Black diamond represents the mean difference across participants. Participants who had extended PBM application time (7 days) are highlighted in red; d) Relative decay rate (k) of phosphocreatine pre- and post-PBM treatment. Small dots represent individual data. Blue triangle: mean k rate pre-PBM. Orange square: mean k rate post-PBM. No significant difference pre and post treatment was found (Wilcoxon Signed Ranks test, two-tailed, Z = −0.338, p = 0.813); e) Relative decay rate (k) of phosphocreatine pre- and post-PBM. Coloured lines indicate individual data. Dashed black line and dots represent the mean of all participants; f) Difference in k-rate (post-PBM – pre-PBM). Small dots indicate individual data. Black diamond represents the mean difference across participants.

To determine whether participant age influenced the change in k-rate of Pi, a Kendall’s tau (non-parametric) correlation was performed between age (years) and the difference in k-rate before and after PBM treatment. The relationship between age and Pi k-rate change was not significant (Kendall’s tau b = −0.293, p = 0.362), indicating that age was not a significant confound within the range tested (60 – 85 years).

To verify that the increase in rate of ATP synthase post-PBM treatment did not simply reflect a whole-body change in metabolism, we applied the same analysis (with *T*_1_ set to 5.1s) to explore creatine kinase-based anaerobic metabolic pathways. Figure 6d-f summarises the changes in the rate of decay of PCr before and after PBM treatment. In contrast to Pi, there was no significant change in decay rates of PCr before and after PBM treatment (Wilcoxon Signed Ranks test, two-tailed, Z = −0.338, p = 0.813).

It is noted that participants undergoing either 4- or 7-day PBM treatment showed significant increases in flux rate. However, there was no apparent association between the length of PBM treatment and *k_f_* rate change across our limited cohort (figure 6c).

By quantifying the chemical shift offset, o_pi_, between Pi(i) and PCr, ^31^P MRS data can also be used to extract intracellular pH information. Using equation 11, ^105^ we estimated pH before and after PBM treatment. While there is a small increase in pH post PBM, 7.54 ± 0.01 compared to 7.50 ± 0.01 pre PBM, this is within standard error and therefore likely not a driver or reflective of the measured increase in ATP synthase flux.

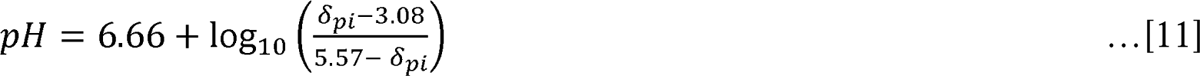

### Light Transport through Brain Tissue

MCS results for grey matter and cytochrome-c absorption are shown in figure 7a-d and 7e-h respectively. Values are given in terms of fractional photon weight (fpw). Note that the black line corresponds to an informed MCS model based on participant MRI data. Segmentation of structural MRI data revealed mean thickness of scalp = 7.4 ± 0.9mm; skull = 4.5 ± 0.6mm; GM = 2.3 ± 0.1mm in-line with published values.^106^ ^107, 108^

**Figure 7:**
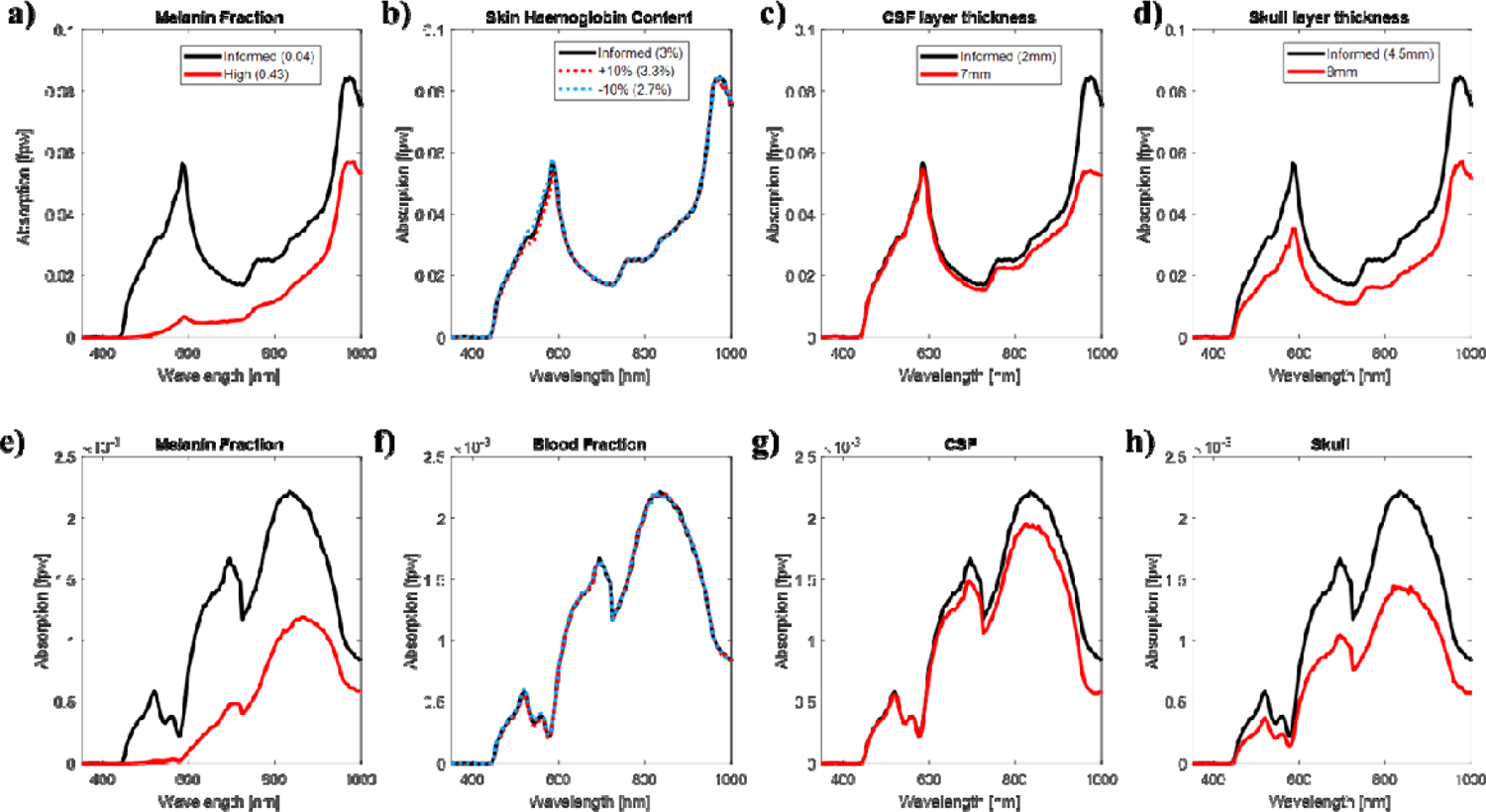
Extensive simulations of absorbed light in the cortical grey matter. Total grey matter light absorption across different physiological parameters including melanin concentration, skin haemoglobin content, CSF and skull thickness are shown in panels a-d respectively. The MRI informed model shows absorption peaks in GM at ∼590 nm and 970 nm. These correspond to absorption by haemoglobin and water respectively. This absorption profile is multiplied by the fractional μ*_A_ of cytochrome-c oxidase in grey matter to explore the equivalent fraction of incident photon weight actually absorbed by cytochrome-c oxidase in the grey matter (panels e-h). There is a peak in photon absorption by cytochrome-c at the illuminated 670 nm point. Data suggest that illumination at 830 nm would result in slightly improved absorption and therefore activation of cytochrome-c oxidase. As we age our skull thickness can vary; using the light transport model we find skull thickness has a quite significant impact on absorption, with the cytochrome-C oxidase absorption being approximately halved in models with thick skull (8 mm) compared to data informed (4.5 mm). We see increased CSF volume with age; increasing the thickness of the CSF layer leads to a slight decrease in total grey matter absorption towards the higher wavelengths, which only slightly decreases cytochrome-c oxidase absorption in the grey matter. Melanin fraction has a marked effect on absorption indicating skin tone is an important consideration for PBM treatment; a change in skin haemoglobin content of ±10% does not significantly change the results, despite the high absorption coefficient*.

Simulations indicate that at any given wavelength (above 450nm) less than 8% of the light transmitted into the head reaches the GM (figure 7a-d). Note that this number does not account for the immediate reflection or refraction of light – just that absorbed. The highest quantity of light traversing to the GM layers corresponds to the high IR wavelength range (∼1000 nm). Due to the high-water content, and hence high absorption in brain tissue, few of these low energy photons will be absorbed by cytochrome-c in the mitochondria. At the target wavelength (670nm) around 2% of light is absorbed in GM.

Elucidating potential PBM mechanisms, we considered absorption of photons by cytochrome-c in the mitochondria in the GM. GM absorption (in fpw) is shown in figure 7a-d. Multiplication of this GM absorption profile by the fractional μ_A_ of cytochrome-c one can explore the fraction of incident photon weight actually absorbed by cytochrome-c oxidase in the GM (figure 7e-h). We provide theoretical evidence that ∼0.16% of 670 nm photons incident on the GM are absorbed by cytochrome-c oxidase. Peak cytochrome absorption occurs between 810-820nm at 0.2%.

The sensitivity of the simulation to different tissue model parameters was tested. Simulations show that across the UV range (350-450 nm) absorption by our skin dominates (data not shown). As is known, our skin layers play an important role in preventing harmful high energy photons from reaching the skull/brain. Mitochondria are therefore semi-protected in everyday life from the potential reductions in intracellular ATP production induced by cytochrome-c absorption at these wavelengths.^63^ These wavelengths are believed to reduce interfacial water layer viscosity and damage mitochondria.^65^ It is the melanin in the skin that dominates absorption. To explore changing melanin fraction in the skin layer from 0.04 (representing pale skin tones) to 0.43 (representing dark skin tones)^109^ on the amount of light being absorbed by GM or cytochrome-c in the brain (figure 7a & 7e respectively). We demonstrate a dramatic 2/3 drop in the amount of 670nm photons being absorbed by cytochrome-c in GM under skin with high melanin fraction. The drop is less pronounced (1/2 photon count) at the optimum 820 nm peak.

Additionally, the sensitivity to skin haemoglobin content was assessed (figure 7b & 7f), as haemoglobin is the dominant absorber in the tissue model over the visible wavelengths. However, we found that a change in skin haemoglobin content of ±10% did not significantly change the results, despite the high absorption coefficient. This probably reflects that at the IR range of wavelengths light travels unperturbed through the superficial dermal skin layers.

CSF thickness varies with the shape of the cortical surface. Sulci depth and width could be affected by age, but as an estimate, large sulci are around 16 mm deep.^99^ The thickness of the CSF volume was inflated from 2 mm to 7 mm to indicate the effect this would have on grey matter absorption, shown in figure 7c & 7g. Simulations confirm that the relative peak height between 670 and 820 nm reduces for thicker CSF layers, but 820 nm remains the optimum light wavelength for cytochrome-c absorption even at this 7 mm extreme.

Sawosz et al. ^98^ states a typical maximum skull thickness is around 8 mm. The skull layer was inflated to match this mean and results in figure 7d & 7h compare a thick 8 mm skull layer with the MRI informed estimate of 4.5 mm. As expected, we see a ∼30% drop in CCO absorption at 670nm behind thicker skull regions.

## Discussion

It has been previously hypothesised across the literature that light activation of the ETC mediator cytochrome-c and the associated protein cytochrome-c-oxidase drive increases in ATP synthase exchange rate. Using magnetisation transfer ^31^P MRS we have quantified a significant increase in neuronal aerobic ATP flux in response to 670 nm PBM treatment over 5 days in a small cohort of aged (60 years+) healthy participants. There is strong evidence for direct glycolysis/mitochondrial involvement as we found no observable increase via the creatine kinase pathway.

While we do observe a significant increase in ATP synthase rate post PBM treatment and such data may lend support to the theory that PBM activates CCO in the ETC, we acknowledge that the number of photons reaching the mitochondria for absorption is extremely small (∼0.15%), as demonstrated by MR informed theoretical light transport model. Wide spectral simulations indeed confirm that 670 nm represents a small peak wavelength for cytochrome c absorption for transcranial PBM however, if cytochrome absorption is the leading mechanism, applying PBM at 810-820nm would improve therapeutic efficacy. In line with the results of Wong-Riley et al. (2004) our simulations show a dip in light absorption at ∼730nm and would indicate this wavelength will be less effective for PBM therapy if cytochrome c absorption is the leading mechanism.

Alternative mechanisms other than cytochrome c absorption are now widely discussed across the literature to explain improved mitochondrial function with PBM.^15, 65^ Moreover, other physiological confounds could be driving the increase in ATP flux rate measured here. Recent studies have demonstrated that PBM directly affects cerebral blood flow, CBF, cerebral oxygenation, ^110^ regulation (via nitric oxide generation),^111^ ^16, 112, 113^ although such effects have been shown to be somewhat wavelength dependent.^114^ Notwithstanding, it is known that with age tissue pO_2_ drops.^115^ Therefore, one could hypothesise that an increase in CBF could drive increases in oxygen extraction fraction under such modulated oxygen concentration gradients (between blood vessels and tissue). This would positively affect cognitive function as others have found.^112^ How long CBF remains elevated after PBM treatment would need to be determined as part of future experiments utilising quantified arterial spin labelling approaches.^116^

Nevertheless, in agreement with previous studies we show that, at suitable wavelengths (red to near infrared), light used for transcranial PBM easily penetrates ∼20-30 mm through the scalp and skull.^15^ However, a wide range of both anatomical and physiological parameters need to be taken into account when considering how much of the incident light can penetrate through to the brain. These parameters include individual head geometry and relative tissue composition which can be dictated to a certain extent by age and gender. Depending on the region of interest of the brain, the scalp-brain distance can vary considerably^117, 118^ and therefore could affect the depth at which light penetration can reach.^119–121^ Skin colour due to the amount of melanin and hair colour (if treatment is not limited to the forehead region) could also play a considerable role as they will absorb the light and attenuate the amount reaching the grey matter. Optical parameters also need to be considered such as scattering coefficient, wavelength, exposure time, treatment area, coherence and pulse structure (continuous wave or pulsed).^122^ Standard structural MRI can help quantify some of the above to better improve light models and better inform PBM treatment regimes. While 3D tissue models informed by anatomical MRI data would be beneficial, computational cost can become prohibitive if cluster computing is not available.

One could assume that no matter the mechanism, strategies to increase the number of incident photons would result in further increases in ATP flux. Other alternatives to transcranial light delivery to overcome the barrier of scalp and skull include intracranial, intranasal^23^ for delivery of the light directly through the oral cavity or through the retina, however all of these are more or less invasive to varying degrees and are dependent on which area of the brain the therapy is needed. Our informed simulations relating to transcranial PBM can be used to choose appropriate wavelengths at the peaks of various tissue absorption to help explore, through quantified MRS the metabolic benefits. Again, if cytochrome-c is involved the optimal illumination wavelength would be 810-20nm (figure 7). This indicates that future PBM studies exploring increasing/restoring brain related ATP synthase should use this higher wavelength of light, under the leading hypothesis that absorption is causing an effect. However, as we near the infrared range, absorption will be affected by the presence of thicker water/CSF layer (as water absorbs high wavelength photons). This finding is in line with Jagdeo et al (2012) who demonstrated using cadaver skull with intact soft tissue, that near infrared light at 830 nm penetrated further than red light at 633 nm (Temporal region 0.9% @830 nm, 0% @633 nm; frontal region 2.1% @830 nm, 0.5% @633 nm; Occipital region 11.7% @830 nm, 0.7% @633 nm).^57^ Smaller relative drops in absorption at 670nm were observed here (figure 7e) and therefore to avoid potential bias due to skull thickness it could prove the better illumination wavelength for PBM. Here our MRI informed simulations could help mitigate the effects of varying skull thickness and allow determination of the best position and angle for placing the bulb to maximise absorption in the GM.

Racial bias in pulse oximetry and functional near-infrared spectroscopy (leading optical techniques relying on blood absorption-based measures) has recently come into focus.^123, 124^ ^125^ Melanin fraction (which determines skin tone) has a marked effect on absorption (figure 7f) and must therefore be an important consideration when designing PBM treatment. These data will help estimate the potential impact of PBM when applied to patients with darker skin tones. Either the incident light will have to be increased or PBM light exposure time increased to overcome this loss. Increasing the energy density of the incident light, while overcoming this potential PBM shortfall, may impact physiology in other unknown ways. Note, the energy required for effective PBM is low, in the range of 1 to 16 joules/cm^2^. It has been shown that above a certain threshold (outside this dose window) increasing the energy will not increase the therapeutic effects.^126^ Our study here delivered the recommended 20 minutes treatment time over the course of several days to reach the optimum dose. Increasing time to overcome higher melanin absorption may become counterproductive. Future experiments should explore this in depth and validate potential impact on ATP synthase with ^31^P MT MRS. We acknowledge that a failure of the current study was that the small all white cohort did not allow us to fully quantify such effects for ^31^P MT MRS metabolic measures.

We note that the small signal amplitude of Pi can limit quantification of ATP synthesis with MT based ^31^P MRS. Preclinical studies with progressive saturation transfer at high magnetic fields (11.7T) easily differentiate/resolve the intra- and extracellular Pi pools.^127^ Tiret et.al. ^127^ noted that failing to accurately resolve extracellular Pi can cause a significant bias in the estimation of the forward constant rate of ATP synthesis. However, in the present study we demonstrate that it is still possible to extract important metabolic parameters even at clinically relevant MRI field strengths (3T) with suitable model fitting and data filtering.

While the exact mechanisms of magnetisation/saturation transfer effects are still debated^73, 128–130^, one could alternatively use inversion transfer (IT) regimes. The main advantages of using IT to quantify exchange kinetics^131, 132^ is that (i) it does not require long saturation pulses; (ii) is less susceptible to unintended MT effects from small pools of metabolites and (iii) the sensitivity to ATP synthesis rate can be enhanced by inverting PCr and all ATP resonances.^133^ In this way the recovery of γ-ATP is significantly delayed and results in amplified MT effects between γ-ATP and Pi. The disadvantage is that more comprehensive modelling is required to fully account for multiple magnetisation exchanges including the cross-relaxation between γ-ATP and β-ATP (i.e. the nuclear Overhauser effect^134^). This method should be tested in response to PBM treatment but would require investment in RF coils capable of transmitting a homogenous 180° B_1_ field and consideration of SAR.

Chaumeil et al. (2009) performed a multimodal imaging study based on the combination of three neuroimaging techniques using ^18^F-FDG PET (glucose consumption), indirect ^13^C MRS (TCA cycle flux) and ^31^P MT-MRS (rate of ATP synthesis). The consistency of these three techniques for measuring metabolic fluxes does demonstrate the robustness of magnetisation transfer ^31^P MRS for directly evaluating ATP synthesis in the living brain across species.^135^–137

The ^31^P MT MRS method (with variable saturation time) deployed here to validate the positive metabolic benefits of PBM could be tested further by investigating exercise related increases in creatine kinase-based metabolism. Following exercise, one should expect to observe a change in ATP k-rate based on the PCr peak changes, without a change in the Pi peak. Future experiments could also compare the change in Pi related ATP production with PBM treatment in a young cohort, as PBM is purported to be most effective in older animals/humans^43^. Finally, research using similar methods in patients with neurodegenerative disease in which mitochondrial function (and hence ATP production) is further impaired. This would help validate that PBM is indeed a useful neuroprotective technique in aging and disease. It has been reported through the NADPARK study^138^ that nicotinamide riboside taken as an oral daily supplement improves mitochondrial function in early-stage PD. Our ^31^P MT MRS method could be used to measure this improvement in terms of ATP synthesis rate and one could envisage using a combined treatment of transcranial photobiomodulation and nicotinamide riboside supplement. We are excited about the future of our refined PBM protocol for use in the fight against neurodegenerative disease.

## Author Contributions

EF: Data curation, Formal Analysis, Funding Acquisition, Investigation, Methodology, Project Administration, Resources, Software, Validation, Visualisation, Writing original draft, Writing review and editing.

FT: Data curation, Formal Analysis, Investigation, Methodology, Project Administration, Resources, Software, Validation, Visualisation, Writing original draft, Writing review and editing.

KC: Data curation, Formal Analysis, Validation,

MS: Resources, Validation, Visualisation, Writing review and editing.

AO: Data curation, Formal Analysis, Methodology, Resources, Software, Validation, Visualisation, Writing review and editing.

GJ: Visualisation, Writing review and editing.

HB: Data curation, Formal Analysis, Investigation, Methodology, Project Administration, Resources, Software, Validation, Visualisation, Writing original draft, Writing review and editing.

AK: Data curation, Formal Analysis, Investigation, Methodology, Project Administration, Resources, Software, Validation, Visualisation, Writing original draft, Writing review and editing.

## Acknowledgements

We thank the staff at the York Neuroimaging Centre for support with 3T MRI scanning and participant handling. Theoretical light transport modelling was undertaken on the Viking Cluster, which is a high-performance computing facility provided by the University of York. We are grateful for computational support from the University of York High Performance Computing service, Viking and the Research Computing team. We thank Dr Alexandra M. Olaru, Siemens Healthineers, Siemens Healthcare Ltd, Surrey, UK, GU16 8QD, for initial support with IDEA programming in VE11C.

## Abbreviations

ADP: adenosine diphosphate

ATP: adenosine triphosphate

BW: bandwidth

Cr: creatine

ETC: electron transport chain

FWHM: Full width at half maximum

Pi: inorganic phosphate

MCS: Monte Carlo simulation

MRS: magnetic resonance spectroscopy

MT: magnetisation transfer

PBM: photobiomodulation

PCr: phosphocreatine

RF: radiofrequency

ROS: reactive oxygen species

TE: Echo time

TR: Repetition time.

